# Rostro-caudal TMS mapping of immediate transcranial evoked potentials reveals a pericentral crescendo-decrescendo pattern

**DOI:** 10.1101/2025.02.14.638272

**Authors:** Marten Nuyts, Mikkel Malling Beck, Agata Banach, Axel Thielscher, Raf Meesen, Leo Tomasevic, Hartwig Roman Siebner, Lasse Christiansen

**Affiliations:** Danish Research Centre for Magnetic Resonance, Department of Radiology and Nuclear Medicine, Copenhagen University Hospital - Amager and Hvidovre, Copenhagen, Denmark, Kettegård Allé 30, 2650 Hvidovre, Denmark; REVAL - Rehabilitation Research Center, Faculty of Rehabilitation Sciences, University of Hasselt, Diepenbeek, Belgium; Department of Health Technology, Technical University of Denmark, Kgs. Lyngby, Denmark; Movement Control and Neuroplasticity Research Group, Department of Movement Sciences, Group Biomedical Sciences, KU Leuven, Leuven, Belgium; Department of Psychiatry and Psychotherapy, University of Regensburg, Regensburg, Germany; Department of Human Sciences, Institute of Psychology, University of the Bundeswehr Munich, Neubiberg, Germany; Department of Neurology, Copenhagen University Hospital - Bispebjerg and Frederiksberg, Bispebjerg Bakke 23, 2400 Kø benhavn NV, Denmark; Department of Clinical Medicine, Faculty of Health and Medical Sciences, University of Copenhagen, Blegdamsvej 3B, 2200 Copenhagen N, Denmark; Institute of Neuroscience, University of Copenhagen, Blegdamsvej 3B, 2200 Copenhagen N, Denmark

**Keywords:** Transcranial magnetic stimulation (TMS), Electroencephalography (EEG), TMS-EEG, immediate transcranial evoked potential (iTEP)

## Abstract

**Background:** We recently demonstrated that single-pulse TMS of the primary sensorimotor hand area (SM1_HAND_) elicits an immediate transcranial evoked potential (iTEP). This iTEP response appears within 2–7 ms post-TMS, featuring high-frequency peaks superimposed on a slow positive wave. Here, we used a linear TMS-EEG mapping approach to characterize the rostro-caudal iTEP expression and compared it to that of motor-evoked potentials (MEPs).

**Methods:** In 15 healthy young volunteers (9 females), we identified the iTEP hotspot in left SM1_HAND_. We applied single biphasic TMS pulses at an intensity of 110% of resting motor threshold over six cortical sites along a rostro-caudal axis (2 cm rostral to 3 cm caudal to the SM1_HAND_ hotspot). We analyzed site-specific iTEP and MEP responses.

**Results:** iTEP magnitude decreased rostrally and caudally from the SM1_HAND_ hotspot. MEPs exhibited a similar rostro-caudal crescendo-decrescendo pattern. While iTEP and MEP response profiles were similar, normalized iTEP amplitudes decayed less rapidly at the first postcentral site.

**Discussion:** These findings support the idea that pericentral iTEPs reflect a direct response signature of the pericentral cortex, possibly involving a synchronized TMS-induced excitation of cortical pyramidal tract neurons. Similar but non-identical rostro-caudal patterns suggest that iTEPs and MEPs may arise from overlapping but distinct neuronal populations.

## 1. Introduction

Transcranial magnetic stimulation (TMS) synchronously excites cortical neurons, and neuronal excitation propagates transsynaptically through cortico-cortical and cortico-subcortical pathways [1]. Simultaneous electroencephalography (EEG) captures these transcranially evoked potentials (TEPs), reflecting the temporospatial dynamics of TMS induced activation of local and remote brain regions [2, 3], offering insights into cortical excitability and connectivity [4, 5]. Measuring the immediate cortical response to single-pulse TMS has been challenging, as the EEG signal within the first 10 milliseconds was obscured by various artifacts generated by the TMS pulse [6, 7] and scalp muscle activity [8].

Using single-pulse TMS over the left primary sensorimotor hand area (SM1_HAND_), we recently demonstrated the feasibility of recording immediate transcranial evoked potentials (iTEPs) as early as 2-3 ms post-stimulation [9]. A subsequent study confirmed the presence of iTEPs using a different TMS-EEG setup [10]. These responses feature high-frequency peaks (650-900Hz inter-peak frequency) superimposed on a slower wave. The short latencies of iTEPs align with evoked neuronal activity recorded invasively in non-human primates [11], possibly reflecting the excitation of large pyramidal neurons. The rhythmicity of the high-frequency iTEP components resembles TMS-evoked descending corticospinal volleys, that can be recorded with epidural electrodes [12]. Notably, iTEP inter-peak intervals mirror those observed when probing short-interval intracortical facilitation (SICF) with paired-pulse TMS of SM1_HAND_ and measuring motor-evoked potentials (MEPs) in the contralateral hand muscles [13]. Establishing iTEPs as a direct marker of cortical excitability enhances the utility of the TMS-EEG approach, as later TEP components likely reflect secondary activation in local and remote brain regions, alongside non-transcranial co-activations via multisensory peripheral inputs [14, 15].

In our initial report, iTEPs were found to depend on TMS intensity and current direction [9]. Moreover, iTEPs could only be elicited from the SM1_HAND_ region and not from posterior parietal or midline areas, despite identical stimulation parameters. While these findings might suggest that iTEPs are a characteristic response feature of the sensorimotor cortex, this remains to be systematically investigated.

This study aimed to map the spatial expression of pericentrally evoked iTEPs along a rostro-caudal gradient. Using the same target region as in previous iTEP studies [9, 10], we stimulated six frontoparietal cortical sites, spaced 1 cm apart, spanning 2 cm rostral to 3 cm caudal from the left SM1_HAND_. Since the neural circuitries generating high-frequency cortical activity (650-900Hz) are thought to reside in the pericentral cortex [16], we hypothesized that iTEPs would peak over SM1_HAND_ and gradually decline with increasing distance in both, the rostral and caudal direction. Concurrently, we recorded MEPs from two contralateral intrinsic hand muscles to examine the spatial alignment and separation of iTEP- and MEP-generating sites with our linear TMS mapping approach.

## 2. Methods

### 2.1. Participants

A total of 17 healthy volunteers (10 females; mean age: 27; range: 20-35) were initially recruited to participate. Participants were excluded in case of (1) contraindications for TMS or MRI; (2) a known neurological disease or psychiatric disorder; (3) coactivation of scalp muscles following a tailored SM1_HAND_ hotspot procedure (see below). Since higher absolute TMS intensities are more prone to coactivate scalp muscles, participants were partly, but not exclusively, recruited based on their resting motor threshold (rMT). It was possible to collect scalp muscle artifact-free data in 15 out of the 17 individuals tested. All participants provided written informed consent prior to participation, following the Declaration of Helsinki. The protocol was approved by the regional ethical committee (Capital Region of Denmark; Protocol number H-15008824).

### 2.2. Experimental design

A tailored SM1_HAND_ hotspot procedure was used (for details, see [9] and **Figure 1**) to allow the recording of the earliest EEG responses following TMS. The left motor hand knob area, identified on the individual’s T1-weighted image, was used as a starting point for stimulation [17]. Starting from the motor hand knob area, the coil was moved in small steps to locate the stimulation site with the largest and most consistent MEP amplitudes in the FDI muscle, designated as the ‘MEP hotspot’ (**Figure 1**). Over the MEP hotspot, we determined the individual resting motor threshold (rMT), defined as the lowest stimulation intensity capable of eliciting peak-to-peak MEP amplitudes of ≥50 µV in at least 5 out of 10 consecutive trials. Next, while stimulating at a suprathreshold intensity of 110%rMT, 10-20 averaged EEG trials were visually checked online for scalp muscle artifacts using the BrainVision Recorder Software [18]. If no scalp muscle artifacts were observed, which was the case for 10 out of 17 subjects, the MEP hotspot was set as the SM1_HAND_ stimulus location. When scalp muscle artifacts were identified, small adjustments in the coil’s position and tilt were made towards the midline in search for an alternate stimulus location, where MEPs could be evoked without concurrent scalp muscle activation [8]. This position was designated as the “TEP-optimized SM1_HAND_ hotspot”. Over the TEP-optimized SM1_HAND_ hotspot, we reassessed the individual rMT and the absence of scalp muscle artifacts using that location’s target intensity of 110%rMT. If no scalp muscle artifacts were observed, the TEP-optimized SM1_HAND_ hotspot was defined as the SM1_HAND_ stimulus location. This was the case for 5 out of 7 remaining subjects. For the remaining 2 participants, we were unable to identify a scalp muscle artifact-free stimulation location.

**Figure 1.**
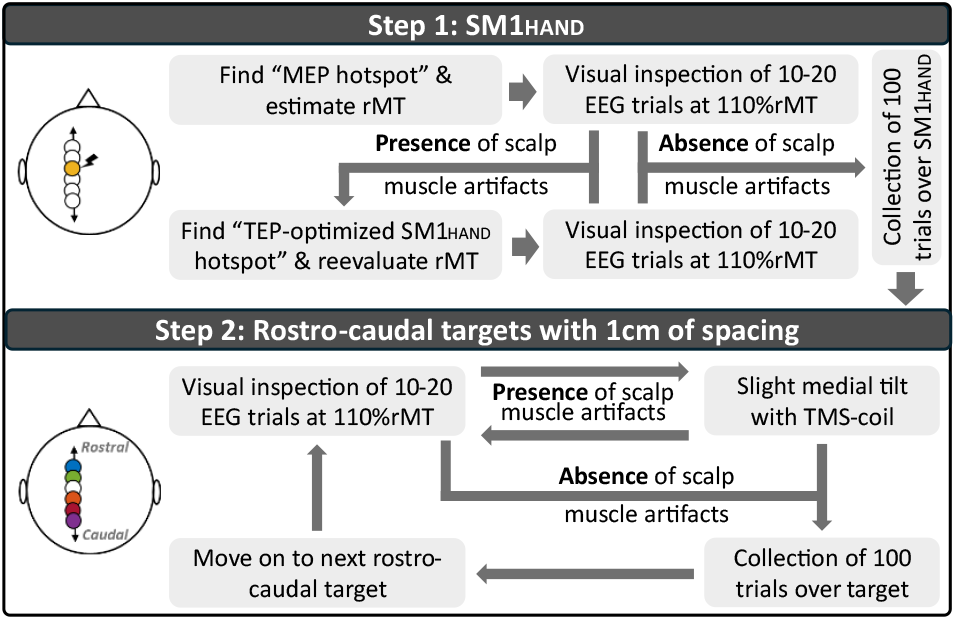
Procedure for collecting a scalp muscle artifact-free dataset at each stimulation site. Step 1 involved identifying the optimal stimulation site where the largest motor-evoked potential (MEP) could be elicited without coactivation of scalp muscles. Small adjustments in coil position and tilt were made if needed. Step 2 involved collecting data along a rostro-caudal line centered around the (TEP-optimized) primary sensorimotor hand area (SM1_HAND_) stimulus location identified in step 1. The coil was tilted slightly medially if scalp muscle coactivation occurred.

In 15 participants (9 females; mean age: 27; range: 20-35), data were collected over six frontoparietal cortical sites along a rostro-caudal line using a stimulation intensity of 110%rMT (**Figure 1**). Each stimulus location was spaced 1 cm apart, moving up to a maximum of 2 cm rostral to 3 cm caudal from SM1_HAND_. We initially set out to also include a stimulation site 3 cm rostral from SM1_HAND_, but according to pilot-testing, this was not feasible due to scalp muscle coactivation by TMS. In line with the procedure outlined for the SM1_HAND_ stimulation site above, each rostro-caudal stimulus location was first visually checked for scalp muscle coactivation (**Figure 1**). If scalp muscle artifacts were observed, small medial coil adjustments were made. This was only the case for two subjects when moving 1cm (SUB-09) and/or 2cm (SUB-02 and SUB-09) rostral from SM1_HAND_. In one participant the rMT value was too high to achieve a stimulation intensity of 110%rMT (SUB-12, rMT = 97%). For this case, the stimulation intensity was set to 100% of the maximum stimulator output.

### 2.3. Transcranial magnetic stimulation (TMS)

Over each stimulus location, single TMS pulses were delivered every 2 s (± 10% jitter) using a 35 mm figure-of-eight coil (MC-B35 coil, MagVenture X100 with MagOption, MagVenture A/S, Farum, Denmark) coated with a layer of foam (± 1.5 mm thick). The coil was placed tangentially to the scalp and oriented approximately 45° to the midline, with the coil handle pointing backwards. A biphasic pulse configuration, inducing an antero-posterior to postero-anterior (AP-PA) current in the brain, was used to lower the absolute TMS intensity required [19, 20]. The recharge delay of the stimulator was set to 500ms post-TMS. Stereotaxic neuronavigation (TMS Navigator, Localite, GmbH, Bonn, Germany) tracked the position, tilt, and orientation of the TMS coil throughout the experiment. In eight participants, structural brain scans were acquired with a 3T PRISMA scanner (Siemens, Erlangen, Germany), while in nine participants a 3T Achieva scanner (Philips, Best, The Netherlands) was used.

### 2.4. Electroencephalography (EEG) and electromyographic (EMG) recordings

EEG was recorded from the scalp with 61 passive Ag/AgCl TMS-compatible electrodes embedded in an equidistant EEG cap (M10 cap layout, BrainCap TMS, Brain Products GmbH, Germany) with the ground and reference electrodes placed on the left and right side of the individual’s forehead, respectively. To keep impedance levels below 5 kΩ, electrodes were prepared with an electroconductive abrasive gel and routinely checked throughout the experiment. Data were recorded with BrainVision Recorder software (Brain Products GmbH, Germany) and a TMS-compatible EEG amplifier (actiCHamp Plus 64 System, Brain Products GmbH, Germany) set at a sampling frequency of 50 kHz (anti-aliasing low-pass filter at 10.300 Hz). A very high sampling frequency was chosen to shorten the duration of the TMS pulse artifact to ± 2 ms [9]. Participants were asked to keep their eyes open, minimize blinking, and relax their face throughout the experiment.

Electromyographic (EMG) activity was simultaneously recorded from the first dorsal interosseus (FDI) and adductor digiti minimi (ADM) muscles of the right hand via self-adhesive surface electrodes (Ambu Neuroline 700) placed on the prepared skin. A belly-tendon montage was used with the ground positioned over the right styloid process of the ulna. EMG signals were amplified (gain = 1000) (Digitimer D360, Digitimer Ltd., Hertfordshire, UK), 0.02 − 2 kHz bandpass filtered, digitized at 5 kHz (CED 1401 micro, CED Limited, Cambridge, UK), acquired via Signal software (Cambridge Electronic Design), and stored for offline analyses.

During blocks of data-collection, pink noise, intermixed with pre-recorded TMS-clicks, was played through modified earplugs to minimize the effect of the TMS-clicking sounds. Sound mask was generated using the TMS Adaptable Auditory Control (TAAC) software [21].

### 2.5. Data analysis

All data were preprocessed and analyzed offline using custom MATLAB scripts (2022b, The Math-Works Inc., Portola Valley, California, USA) using EEGLAB [22] and Fieldtrip functions [23].

EMG data were visually checked for background muscle activity in a window of 100 ms pre-stimulus. Single trials containing background EMG activity were discarded prior to averaging from the EMG and EEG dataset (median: 1; range: 0-28), as were bad channels in the EEG dataset (see **Table S1** for an overview). Peak-to-peak MEP amplitudes from the right FDI muscle were calculated over each stimulus location as the difference between the maximum and minimum values from 20 ms to 40 ms post-stimulus from the average trace.

EEG data were epoched from −500 ms to 500 ms around the stimulus, baseline corrected from a window of −110 ms to −10 ms from the stimulus, band-pass filtered (2nd order zero-phase Butterworth filter at 0.1–2000 Hz), and averaged. Following these steps, the global mean field power (GMFP) was computed [24], as it serves as a general activation index, aimed at avoiding potential spatial bias from single-channel(s) extraction related to the stimulus location. The GMFP was computed with decay artifact channels included and removed. To characterize the high-frequency component of iTEPs, we calculated the difference between the first two peaks relative to the first trough and the third peak relative to the second trough (**Table S2**). Peaks and troughs were calculated over each stimulation location as the local minimum and maxima values from 2 ms to 8 ms post-stimulus from the average trace, with a minimal peak or trough interval of 1 ms.

### 2.6. Statistical analysis

Statistical analysis of all data was performed in MATLAB and R, using RStudio (lme4 package) [25-28]. For simplification, intercepts and error terms are not explicitly reported here but were included in all models. Alpha was set at 0.05, and all P-values were two-tailed. When relevant, Tukey-corrected post-hoc comparisons were performed.

#### 2.6.1. The effect of stimulus location on iTEPs

The effect of stimulus location on iTEPs was examined by applying a temporal clustering approach, with a mixed effect model fitted for each timepoint within a window of 2-8 ms post-stimulation [29]. The mixed models included STIMULATION SITE (six levels, six frontoparietal locations) as a fixed factor, PARTICIPANT as the random intercept, and GMFP as the dependent variable:

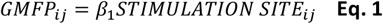

F-values were extracted for each model and used to calculate threshold-free clustering enhanced (TFCE) values [30]:

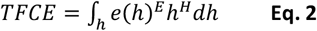

Where *h* represents cluster height (F-value threshold), *e* denotes cluster extent (number of temporally adjacent data points > *h*), and *H* and *E* are their respective weights, with default values of *H* = 2 and *E* = 0.5. *h* was incrementally increased from 0 in steps of 0.2 until the maximum F-value was reached. Significance was derived if TFCE values surpassed the 95th percentile of a surrogate null distribution generated from 800 permutations [29]. By using TFCE, we identified significant differences in iTEPs across stimulus locations over time without performing independent tests at predefined time points.

#### 2.6.2. The relationship between iTEPs and MEPs

The relationship between MEPs and iTEPs was assessed using three complementary analyses. All analyses included normalization procedures to account for individual and measurement variability and aid comparisons between excitability measures.

In the first analysis, we normalized rostro-caudal excitability profiles to each individual’s peak response. Specifically, for each individual excitability measure (MEP, iTEP peak 1 – trough 1, and iTEP peak 2 – trough 1), the amplitude at each location was normalized by dividing it by the amplitude at the stimulation site with the highest response for that individual. Next, a mixed model was derived with AMPLITUDE as the dependent variable and PARTICIPANT included as the random intercept. STIMULATION SITE (six levels, six frontoparietal sites) and EXCITABILITY MEASURE (three levels: MEP; iTEP peak 1 – trough 1; iTEP peak 2 – trough 1) were added as main effects and in interaction with each other:

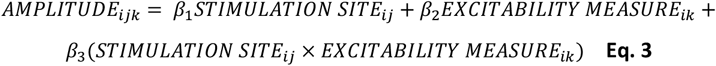

We did not include iTEP peak 3 - trough 2 as a level to EXCITABILITY MEASURE, as this metric could only be extracted in 4 out of 15 participants (see results below).

In the second analysis, we computed an amplitude-weighted mean position for each excitability measure along the rostro-caudal line for each subject [31], designated as the weighted mean position (WMP):

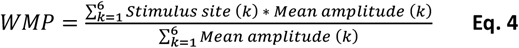

Where “Stimulation site (k)” refers to each of the six stimulation sites, while “Mean amplitude (k)” is calculated from the peak-to-peak amplitudes of MEPs or peak-to-trough amplitudes of iTEPs at each of the six stimulation sites. In doing so, the WMP was calculated for three excitability measures: MEP, iTEP peak 1 – trough 1, and iTEP peak 2 – trough 1. To investigate differences in WMP between MEPs and iTEPs, a mixed model was derived with WMP as dependent variable, PARTICIPANT as the random intercept and EXCITABILITY MEASURE (three levels: MEP; iTEP peak 1 – trough 1; iTEP peak 2 – trough 1) added as a fixed factor:

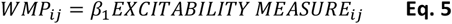

Lastly, the association between the WMP of MEPs and iTEP peaks was tested using a multiple linear regression model:

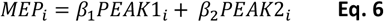

## 3. Results

Single-pulse stimulation over six frontoparietal cortical sites (**Figure 2A**) resulted in MEPs in both hand muscles and prototypical TEPs peaks by stimulating in the proximity of the hand region (**Figure S1**). The iTEPs were evoked in all 15 subjects (**Figure S2**), characterized by a series of 2-3 peaks with an underlying slower component. **Figure 2** illustrates the averaged response for iTEPs (**Figure 2B and 2C**), their averaged topographical distribution between the first and second iTEP peak (**Figure 2D**), and the averaged MEPs in both hand muscles (**Figure 2E**). In addition to **Figure 2**, we recomputed the averaged iTEP response with decay artifact channels removed as shown in the supplementary materials (**Figure S3**).

**Figure 2.**
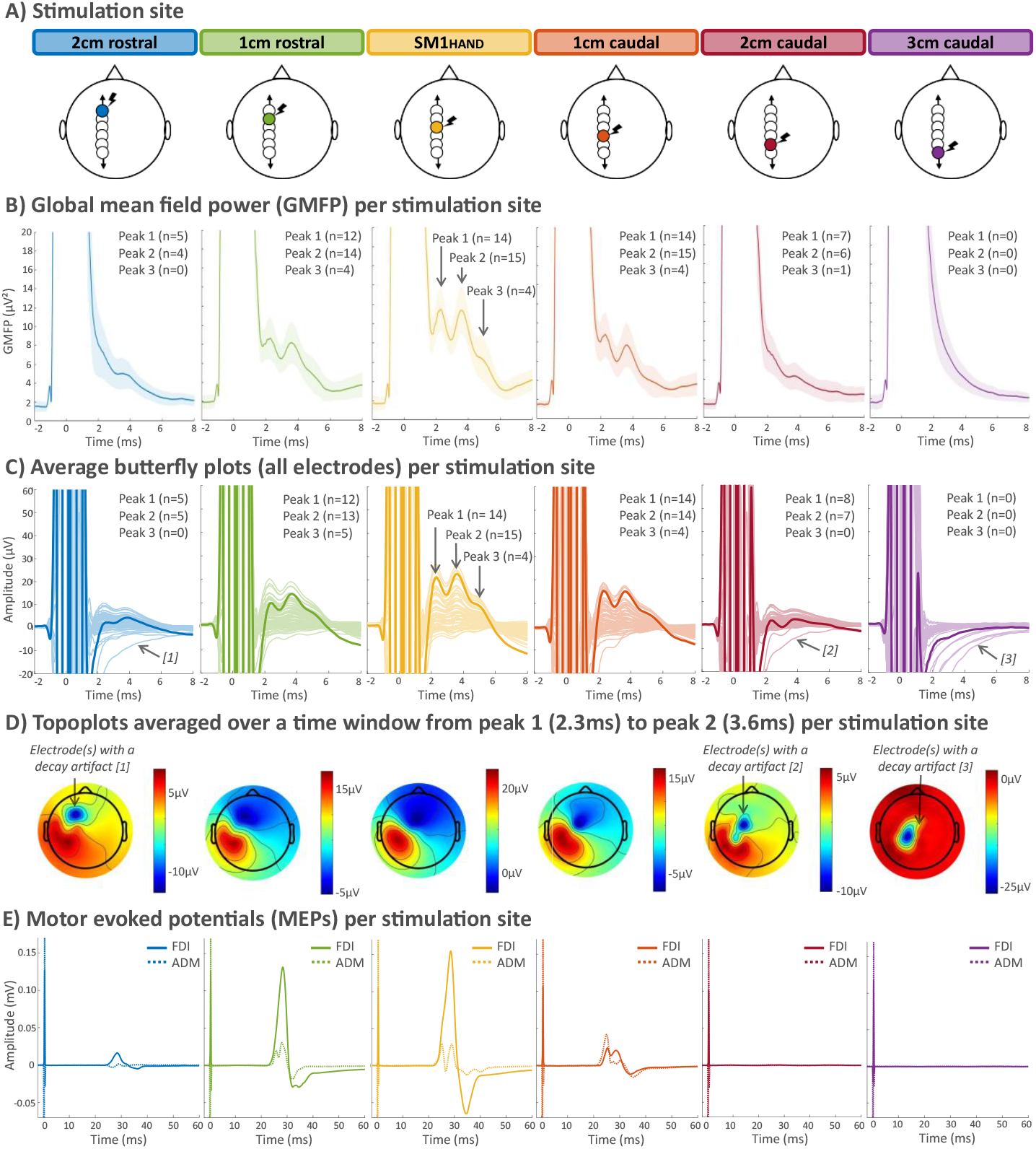
Immediate transcranial evoked potentials (iTEPs) and motor-evoked potentials (MEPs) across stimulation sites. **(A)** The six rostro-caudal sites of stimulation centered around the sensorimotor hand area (SM1_HAND_). **(B)** Global mean field power (GMFP), showing that iTEPs are maximally expressed following stimulation over pericentral stimulation sites. **(C)** Average butterfly plots with the electrode closest to SM1_HAND_ highlighted, showing that iTEPs are maximally expressed at pericentral stimulation sites. Decay artifact channels are indicated by arrows. **(D)** Topoplots, averaged over a time window from iTEP peak 1 to peak 2 (i.e., 2.3ms − 3.6ms). Decay artifact channels are indicated by arrows. **(E)** MEPs of the right first dorsal interosseus (FDI) and abductor digiti minimi (ADM) indicated by solid and dashed lines, respectively. MEP peak-to-peak amplitudes were greatest following stimulation over central sites.

### 3.1. The effect of stimulus location on iTEPs

Decay artifacts were present in some EEG channels after stimulation (**Figure 2C**), and this contaminated the group average EEG data and the topographical distribution of the EEG response (**Figure 2D**). Using a stepwise procedure, we identified and removed channels exhibiting a decay artifact for longer than 2 ms post-stimulation (see **Figure S4** for details). Accordingly, the effect of stimulus location on iTEPs was tested using two separate temporal clustering analyses, with and without decay artifact channels included. Both analyses provided statistical support for previous observations of changes in iTEPs across stimulus locations.

A first temporal clustering analysis, with decay artifacts channels included, revealed a significant difference between stimulation sites driven by a cluster from 3.22 ms to 4.42 ms post-TMS (**Figure 3A**). The median GMFP amplitude was highest when stimulating over SM1_HAND_ (10.16 µV^2^), followed by 1 cm rostral (7.54 µV^2^), 1 cm caudal (6.90 µV^2^), 2 cm rostral (4.88 µV^2^), 3 cm caudal (4.34 µV^2^), and finally 2 cm caudal (4.22 µV^2^) from SM1_HAND_. The same temporal clustering analysis, but with decay artifact channels removed, revealed a significant difference between stimulation sites driven by a cluster from 2.18 ms to 4.42 ms post-TMS (**Figure 3B**). The median GMFP amplitude was when the coil was placed over SM1_HAND_ (9.67 µV^2^), followed by 1 cm rostral (7.31 µV^2^), 1 cm caudal (6.76 µV^2^), 2 cm rostral (4.20 µV^2^), 2 cm caudal (3.71 µV^2^), and finally 3 cm caudal (2.33 µV^2^) from SM1_HAND_.

**Figure 3.**
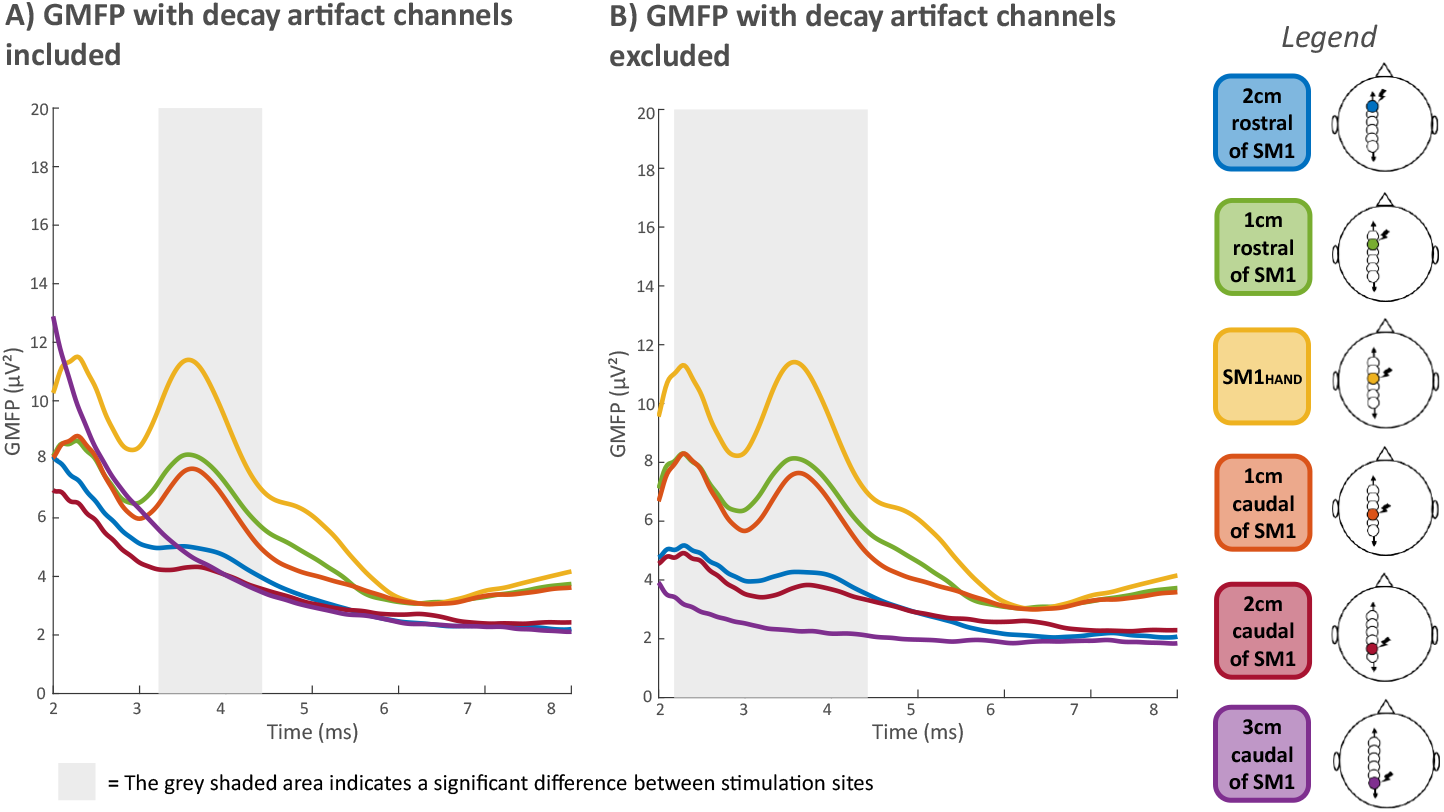
The effect of stimulation site on immediate transcranial evoked potentials (iTEPs). The grey shaded area indicates a significant difference in global mean field power (GMFP) between stimulation sites. **(A)** GMFP with decay artifact channels included. **(B)** GMFP with decay artifact channels excluded.

### 3.2. The relationship between iTEPs and MEPs

Since decay artifacts can (1) obscure the detection of iTEP peaks (**Figure S4**) and (2) alter the time window for detecting effects (**Figure 3**), all analyses examining the relationship between iTEPs and MEPs were conducted using the decay-artifact free data. Overall, we statistically confirm a spatial relationship between MEPs and iTEP peaks, while identifying several important differences.

First, iTEPs and MEPs were elicited concurrently over most (central) stimulus locations, although iTEPs could be elicited in the absence of MEPs during rest (**Figure 4A**). Second, **Figure 4B** illustrates the normalized excitability profiles for both iTEPs and MEPs. A mixed model investigating the effect of stimulation site on these rostro-caudal excitability profiles revealed a significant two-way interaction between STIMULATION SITE and EXCITABILITY MEASURE (F_10,238_ = 2.39, p = 0.01). Specifically, post-hoc testing revealed significant differences between the normalized MEP amplitude and both iTEP peak 1 – trough 1 (*t* = −4.91, p < 0.001) and iTEP peak 2 – trough 1 (*t* = −4.37, p = 0.002) when the coil was positioned 1 cm caudal from SM1_HAND_, as can be observed in **Figure 4B**. In accordance with this result, we observed a median caudal difference of 1.4 mm in the WMP from the first iTEP peak relative to the MEP (**Figure 4C**), though the effect of STIMULATION SITE on WMP did not reach statistical significance (F_2,28_ = 3.02, p = 0.065). Nevertheless, a numerical indication was prevalent, with 12 out of 15 subjects showing a more caudal WMP of the first iTEP peak relative to the MEP. Additionally, an overall moderate linear relationship was observed between the WMP of MEPs and iTEP peaks (**Figure 4**), with an R^2^ of 0.54. Collectively, these results suggest that iTEPs expression resembled that of the MEP, albeit slightly caudal.

**Figure 4.**
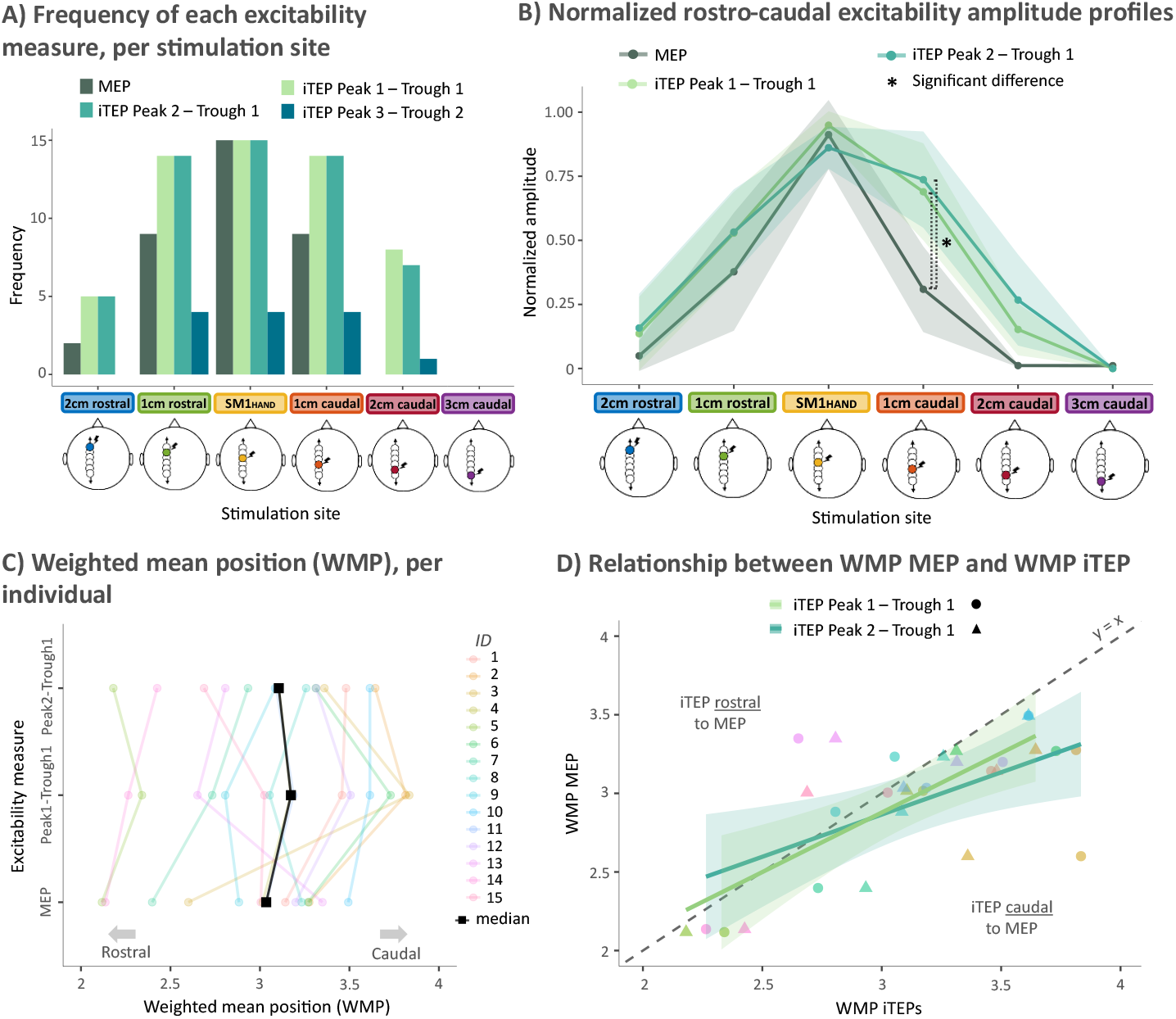
Relationship between the amplitude of immediate transcranial evoked potentials (iTEPs) and motor-evoked potentials (MEPs). **(A)** Histograms showing the presence of MEPs and iTEP responses at each stimulation site across subjects. **(B)** Normalized rostro-caudal excitability amplitude profiles, illustrating the significant EXCITABILITY MEASURE by STIMULATION SITE interaction. **(C)** Weighted mean position (WMP) for each excitability measure shown per subject, with black squares indicating the median WMP for each measure. **(D)** Association between the WMP of MEPs and each iTEP peak, showing a strong, but slight caudal, relationship.

Exploratory analyses revealed a systematic median increase in the onset latency of the MEP and the peak latency of the late iTEP peaks as the coil was centered away from SM1_HAND_ (**Figure S5** and **Table S3**). The analyses and results are described and discussed in the supplementary materials.

## 4. Discussion

This is the first systematic examination of rostro-caudal iTEP expression evoked with single-pulse TMS of the left pericentral hand region. In agreement with our hypothesis, our linear TMS-EEG mapping approach revealed that iTEPs peaked at the SM1_HAND_ hotspot and diminished rostrally and caudally with increasing distance. While rostro-caudal representation of iTEPs and MEPs were closely correlated across stimulation sites, normalized iTEP amplitudes decayed less rapidly than normalized MEP amplitudes in the caudal direction.

### 4.1. The iTEP response peaks at pericentral stimulation sites

The iTEPs diminished with increasing distance from the SM1_HAND_ hotspot. This supports the notion that, given the current stimulation intensity and coil orientation, the cortical circuitries capable of generating the characteristic high-frequency iTEP response pattern are located within the pericentral area [16]. In accordance with our initial report, the temporal iTEP features resemble the rhythmicity of indirect descending corticospinal waves (I-waves) [9]. Findings in non-human primates suggest that stimulation of both premotor and somatosensory areas can evoke I-waves via afferent input to the primary motor cortex [32-35]. Moreover, both electrical stimulation of a peripheral nerve and tactile stimulation of glabrous skin evoke high-frequency activity in S1 in human and non-human primates [36, 37]. Accordingly, iTEPs could still be evoked by stimulation over premotor (i.e., rostral) and somatosensory areas (i.e., caudal) in the present study, but this could also be attributed to spatial spread of the electrical field to the primary (sensori)motor hand area.

### 4.2. Rostro-caudal cortical iTEP and MEP maps are similar but not identical

Our results, showing similar rostro-caudal excitability profiles of iTEPs and MEPs, support the notion that iTEPs reflect a cortical signature of TMS-evoked corticospinal volleys. The inter-individual association between the WMP derived from the iTEP and MEP excitability profiles further suggests neighboring or shared neural generators.

We also observed a slight caudal difference in the local iTEP peak maximum relative to the MEP due to a slower postcentral decay of iTEPs relative to MEPs. This finding suggests a postcentral expansion of the cortical representation of iTEPs relative to the corticospinal representation probed with MEP measurements, as normalized iTEP amplitudes decay less rapidly than normalized MEP amplitudes in the caudal direction. This finding extends our recent finding obtained with sulcus-aligned corticomotor mapping of the intrinsic hand muscles, involving MEP measures of SICF [38]. Sulcus-aligned corticomotor mapping revealed a caudal shift in the WMP of motor maps when generated with SICF adjusted paired-pulse relative to single-pulse TMS [38]. Together, these results suggest that single-pulse MEPs and multi-peak responses, evoked using single-pulse iTEPs and paired-pulse SICF, are generated by overlapping but distinct neural populations.

Unlike SICF paired-pulse protocols, which rely on a subthreshold pulse acting on a suprathreshold one, iTEPs can be elicited in the absence of MEPs. At more rostral or caudal sites, we were still able to observe iTEPs in the absence of MEPs in several participants. This highlights the potential of iTEPs as a direct marker of the immediate motor-cortical response to TMS, including the investigation of sub-motor threshold interactions in the pericentral cortex.

### 4.3. Methodological considerations and recommendations for future work

Our experimental design only allows a rough localization of the maximal iTEP expression across stimulation sites, because we did not systematically optimize TMS parameters such as coil orientation and stimulus intensity, across all stimulus locations. Future studies may benefit from a more fine-grained systematic examination of how specific TMS parameters impact the regional iTEP expression at specific stimulus locations [18]. For instance, we cannot rule out that the absence of MEPs or iTEPs over more rostral and caudal stimulus locations resulted from an insufficient stimulus intensity, a suboptimal pulse configuration or current direction. It would also be worthwhile in future studies to calculate the spatial distributions of the TMS-induced electrical fields and relate them with the iTEP magnitudes [39].

Future studies need to further optimize the avoidance or suppression of decay artifacts, particularly when studying responses at early latencies. The current study showed that these artifacts significantly impact the window to detecting results. Hence, considering hardware specifications, maintaining low impedances, and aligning the electrode lead orientation relative to the TMS coil may be important to mitigate their impact in future research [3].

## 5. Conclusion

Linear rostro-caudal mapping revealed that the characteristic iTEPs peak at the SM1_HAND_ hotspot and decrease rostrally and caudally with increasing distance. The MEPs of contralateral hand muscles exhibit a similar crescendo-decrescendo pattern. Together, these results support the hypothesis that iTEPs reflect a direct response signature of the pericentral cortex, possibly involving a synchronized excitation of pyramidal tract neurons. The rostro-caudal iTEP and MEP maps were very similar but not identical. At the first postcentral site, normalized iTEPs showed a stronger magnitude relative to the normalized MEPs, suggesting that pericentral iTEPs and MEPs may be generated by overlapping but distinct neuron populations.

## Acknowledgments

We would like to thank Sybren Van Hoornweder for the insightful discussions and acknowledge Ditte Haagerup and Kora Montemagno for their support in setting up the experiment.

## CrediT Author Contribution Statement

Conceptualization: MN, MBB, HRS, LC. Methodology: MN, MBB, HRS, LC. Software: MN, MBB, LT, LC. Validation: MN, MBB, LT, HRS, LC. Formal analysis: MN, MBB, LT. Investigation: MN, MBB, AB, LC. Resources: MBB, RM, HRS, LC. Data curation: MN. Writing – Original Draft: MN. Writing – Review & Editing: MN, MBB, AT, RM, LT, HRS, LC. Visualization: MN, MBB, LT, HRS, LC. Supervision: RM, HRS, LC. Project administration: MN, MBB, HRS, LC. Funding acquisition: MN, RM, HRS, LC.

