## Supplementary Materials for "Rostro-caudal TMS mapping of immediate transcranial evoked potentials reveals a pericentral crescendo-decrescendo pattern"

* Shared authorship

### Supplementary tables and figures

Here, we provide supplementary tables and figures to support the methods and results section of the main manuscript.

### Overview of the included number of trials and channels

**Table S1** provides an overview of the total number of trials included and bad channels excluded per subject across stimulation sites.

| SUBJECT | 2cm rostral | | 1cm rostral | | $SM1_{HAND}$ | | 1cm caudal | | 2cm caudal | | 3cm caudal | |
|---|---|---|---|---|---|---|---|---|---|---|---|---|
| | Trials | Channels removed | Trials | Channels removed | Trials | Channels removed | Trials | Channels removed | Trials | Channels removed | Trials | Channels removed |
| SUB-01 | 96 | 0 | 93 | 1 | 97 | 1 | 99 | 1 | 85 | 0 | 97 | 0 |
| SUB-02 | 72 | 3 | 77 | 2 | 74 | 0 | 78 | 0 | 77 | 0 | 72 | 1 |
| SUB-03 | 100 | 3 | 100 | 1 | 100 | 1 | 100 | 1 | 100 | 4 | 98 | 5 |
| SUB-04 | 100 | 0 | 100 | 0 | 98 | 0 | 97 | 0 | 100 | 0 | 100 | 0 |
| SUB-05 | 98 | 3 | 95 | 4 | 94 | 3 | 99 | 3 | 93 | 3 | 94 | 3 |
| SUB-06 | 100 | 3 | 99 | 2 | 100 | 3 | 100 | 2 | 95 | 2 | 100 | 2 |
| SUB-07 | 94 | 1 | 99 | 1 | 96 | 1 | 96 | 1 | 96 | 2 | 98 | 1 |
| SUB-08 | 82 | 0 | 100 | 0 | 89 | 0 | 89 | 0 | 100 | 0 | 94 | 0 |
| SUB-09 | 87 | 2 | 90 | 1 | 100 | 2 | 94 | 2 | 96 | 2 | 92 | 3 |
| SUB-10 | 100 | 2 | 100 | 2 | 99 | 1 | 100 | 1 | 100 | 1 | 100 | 3 |
| SUB-11 | 100 | 1 | 100 | 1 | 100 | 1 | 100 | 2 | 100 | 1 | 100 | 1 |
| SUB-12 | 90 | 4 | 88 | 3 | 99 | 3 | 96 | 2 | 85 | 4 | 93 | 3 |
| SUB-13 | 100 | 0 | 100 | 0 | 100 | 0 | 99 | 0 | 100 | 0 | 81 | 0 |
| SUB-14 | 98 | 0 | 100 | 0 | 100 | 0 | 96 | 0 | 99 | 0 | 100 | 0 |
| SUB-15 | 100 | 1 | 100 | 1 | 100 | 0 | 100 | 0 | 100 | 1 | 100 | 0 |
| Median | 98 | 1 | 100 | 1 | 99 | 1 | 99 | 1 | 99 | 1 | 98 | 1 |

### Overview of the extracted peak-to-troughs of immediate transcranial evoked potentials (iTEPs)

**Table S2** provides an overview of the extracted peak-to-trough amplitudes from the global mean field power (GMFP) for iTEPs. This was done for data with decay artifact channels included and removed. Please note that iTEPs could not be extracted when the coil was moved 3cm caudal from the sensorimotor hand area ($SM1_{HAND}$), as they were absent.

| Decay artifact channels | Stimulus location | Peak-to-trough | Mean | SD |
|---|---|---|---|---|
| INCLUDED | 2cm rostral | Peak 1 – Trough 1 | 2.7 | 2.19 |
| | | Peak 2 – Trough 1 | 1.17 | 0.58 |

| | | | | |
|---|---|---|---|---|
| | | Peak 3 – Trough 2 | / | / |
| | 1cm rostral | Peak 1 – Trough 1 | 2.55 | 2.28 |
| | | Peak 2 – Trough 1 | 2.44 | 2.08 |
| | | Peak 3 – Trough 2 | 1.17 | 0.74 |
| | $SM1_{HAND}$ | Peak 1 – Trough 1 | 3.54 | 2.00 |
| | | Peak 2 – Trough 1 | 3.39 | 2.24 |
| | | Peak 3 – Trough 2 | 1.07 | 0.49 |
| | 1cm caudal | Peak 1 – Trough 1 | 3.24 | 1.64 |
| | | Peak 2 – Trough 1 | 2.51 | 1.96 |
| | | Peak 3 – Trough 2 | 1.48 | 1.88 |
| | 2cm caudal | Peak 1 – Trough 1 | 2.6 | 1.76 |
| | | Peak 2 – Trough 1 | 1.45 | 1.02 |
| | | Peak 3 – Trough 2 | 1.05 | / |
| EXCLUDED | 2cm rostral | Peak 1 – Trough 1 | 2.64 | 2.17 |
| | | Peak 2 – Trough 1 | 1.78 | 1.08 |
| | | Peak 3 – Trough 2 | / | / |
| | 1cm rostral | Peak 1 – Trough 1 | 2.74 | 2.65 |
| | | Peak 2 – Trough 1 | 2.97 | 2.56 |
| | | Peak 3 – Trough 2 | 1.20 | 0.73 |
| | $SM1_{HAND}$ | Peak 1 – Trough 1 | 3.31 | 2.04 |
| | | Peak 2 – Trough 1 | 3.51 | 2.29 |
| | | Peak 3 – Trough 2 | 1.09 | 0.46 |
| | 1cm caudal | Peak 1 – Trough 1 | 3.18 | 1.67 |
| | | Peak 2 – Trough 1 | 2.68 | 2.01 |
| | | Peak 3 – Trough 2 | 1.15 | 1.68 |
| | 2cm caudal | Peak 1 – Trough 1 | 2.27 | 1.64 |
| | | Peak 2 – Trough 1 | 1.57 | 0.99 |
| | | Peak 3 – Trough 2 | 1.05 | / |

**Overview onset latencies across stimulation sites**

**Table S3** provides an overview of adjusted onset latencies for each iTEP peak and the MEP, across each stimulation site. Latencies were adjusted to the most central stimulus location. Please note that iTEP latencies were not extracted when the coil was moved 3cm caudal from $SM1_{HAND}$, as iTEPs were absent over this stimulus location.

| Excitability measure | Stimulus location | Median | SD |
|---|---|---|---|
| iTEP Peak 1 | 2cm rostral | 0 | 0.36 |

| | | | |
|---|---|---|---|
| | 1cm rostral | 0 | 0.01 |
| | $SM1_{HAND}$ | Latencies were adjusted to $SM1_{HAND}$ | |
| | 1cm caudal | 0.04 | 0.39 |
| | 2cm caudal | 0.04 | 0.15 |
| iTEP Peak 2 | 2cm rostral | 0.3 | 0.26 |
| | 1cm rostral | 0.1 | 0.16 |
| | $SM1_{HAND}$ | Latencies were adjusted to $SM1_{HAND}$ | |
| | 1cm caudal | 0.08 | 0.20 |
| | 2cm caudal | 0.38 | 0.26 |
| iTEP Peak 3 | 2cm rostral | / | / |
| | 1cm rostral | 0.2 | 0.3 |
| | $SM1_{HAND}$ | Latencies were adjusted to $SM1_{HAND}$ | |
| | 1cm caudal | 0.33 | 0.38 |
| | 2cm caudal | 0.84 | / |
| MEP | 2cm rostral | 0.3 | 0.14 |
| | 1cm rostral | 0.6 | 1.22 |
| | $SM1_{HAND}$ | Latencies were adjusted to $SM1_{HAND}$ | |
| | 1cm caudal | 0.3 | 0.37 |
| | 2cm caudal | / | / |

### Prototypical TEP peaks across stimulation sites

**Figure S1** illustrates how prototypical TEP peaks, such as N15, P30, N45, P60 and N100, are elicited over $SM1_{HAND}$ and expressed over the remaining stimulates locations.

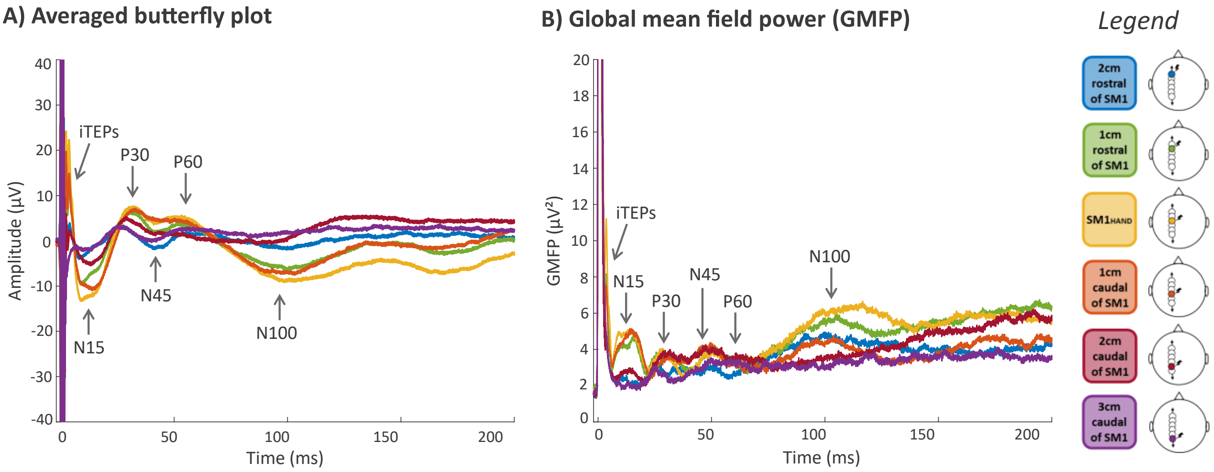

**Figure S1**. Prototypical transcranially evoked potentials (TEP) peaks. **(A)** Averaged butterfly plots, with the electrode closest to the sensorimotor hand area ($SM1_{HAND}$) plotted. **(B)** Global mean field power (GMFP).

### Single-subject iTEPs across stimulation sites

**Figure S2** provides an overview of iTEPs per subject across all stimulation sites. The electrode plotted is the electrode closest to $SM1_{HAND}$, which sometimes suffers from a decay artifact (e.g., SUB-13).

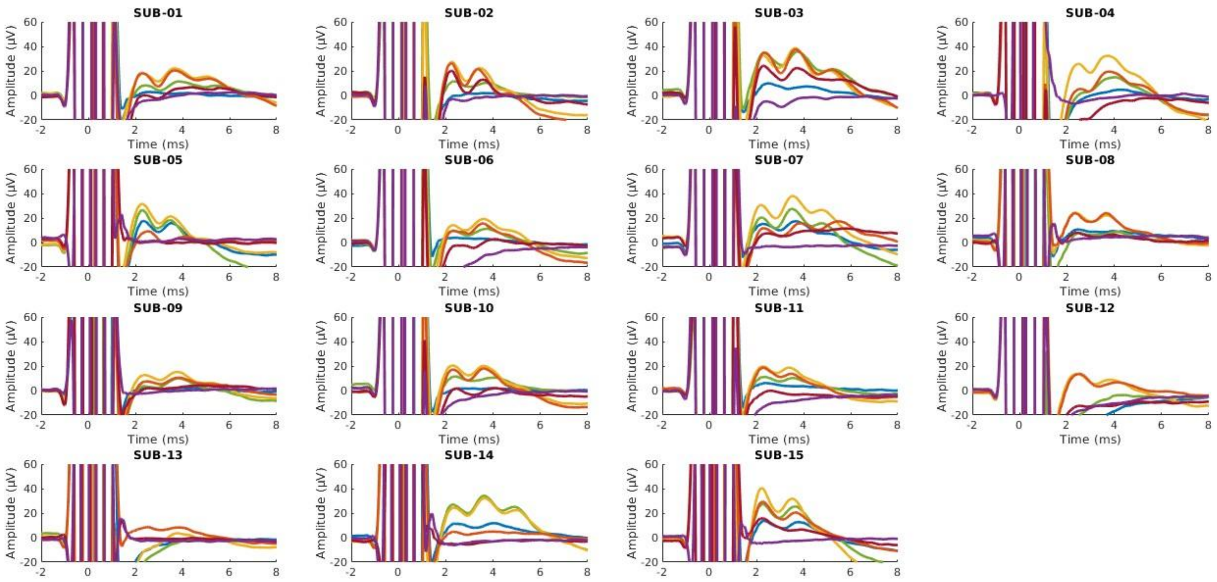


**Figure S2.** Butterfly plots for each individual subject. The electrode plotted is the one closest to the sensorimotor hand area ($SM1_{HAND}$).

### iTEPs with decay artifact channels removed from the dataset

**Figure S3** illustrates iTEPs across all stimulus locations, with decay artifact channels removed from the data. Please note that the topographical plots are min-max scaled per stimulation site.

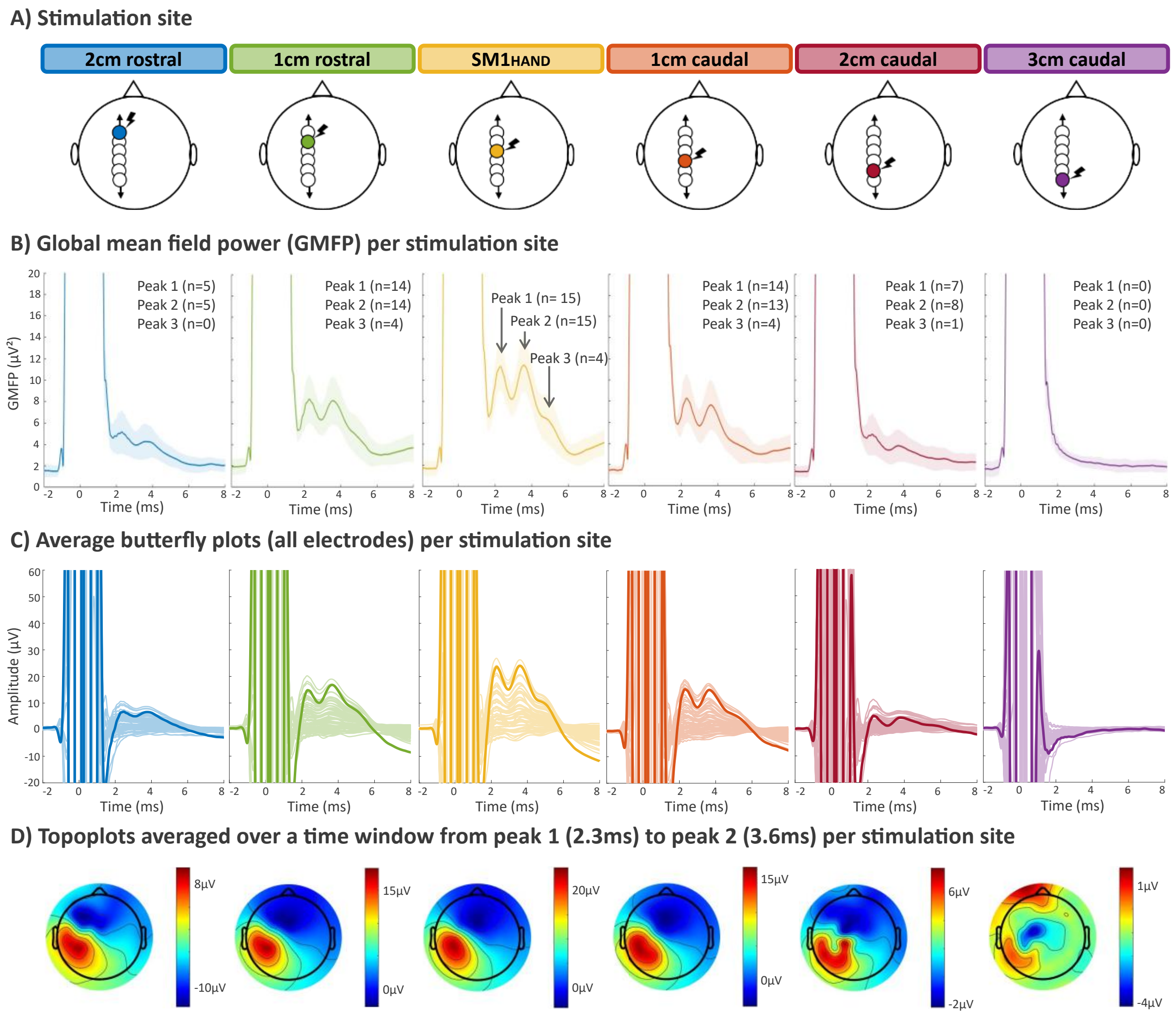


**Figure S3**. Immediate transcranial evoked potentials (iTEPs) with decay artifact channels removed. **(A)** The six rostro-caudal sites of stimulation centered around the sensorimotor hand area ($SM1_{HAND}$). **(B)** Global mean field power (GMFP), showing that iTEPs are maximally expressed following stimulation over pericentral stimulation sites. **(C)** Average butterfly plots with the electrode closest to $SM1_{HAND}$ highlighted, showing that iTEPs are maximally expressed at pericentral stimulation sites. **(D)** Topoplots, averaged over a time window from iTEP peak 1 to peak 2 (i.e., 2.3ms – 3.6ms).

### Decay artifact channels removal

**Figure S4A** shows the challenges posed by decay artifacts as illustrated in an exemplary subject. **Figure S4B** shows the stepwise procedure for identifying and removing channels suffering from a decay artifact and the consequent effects on the GMFP in an exemplary subject.

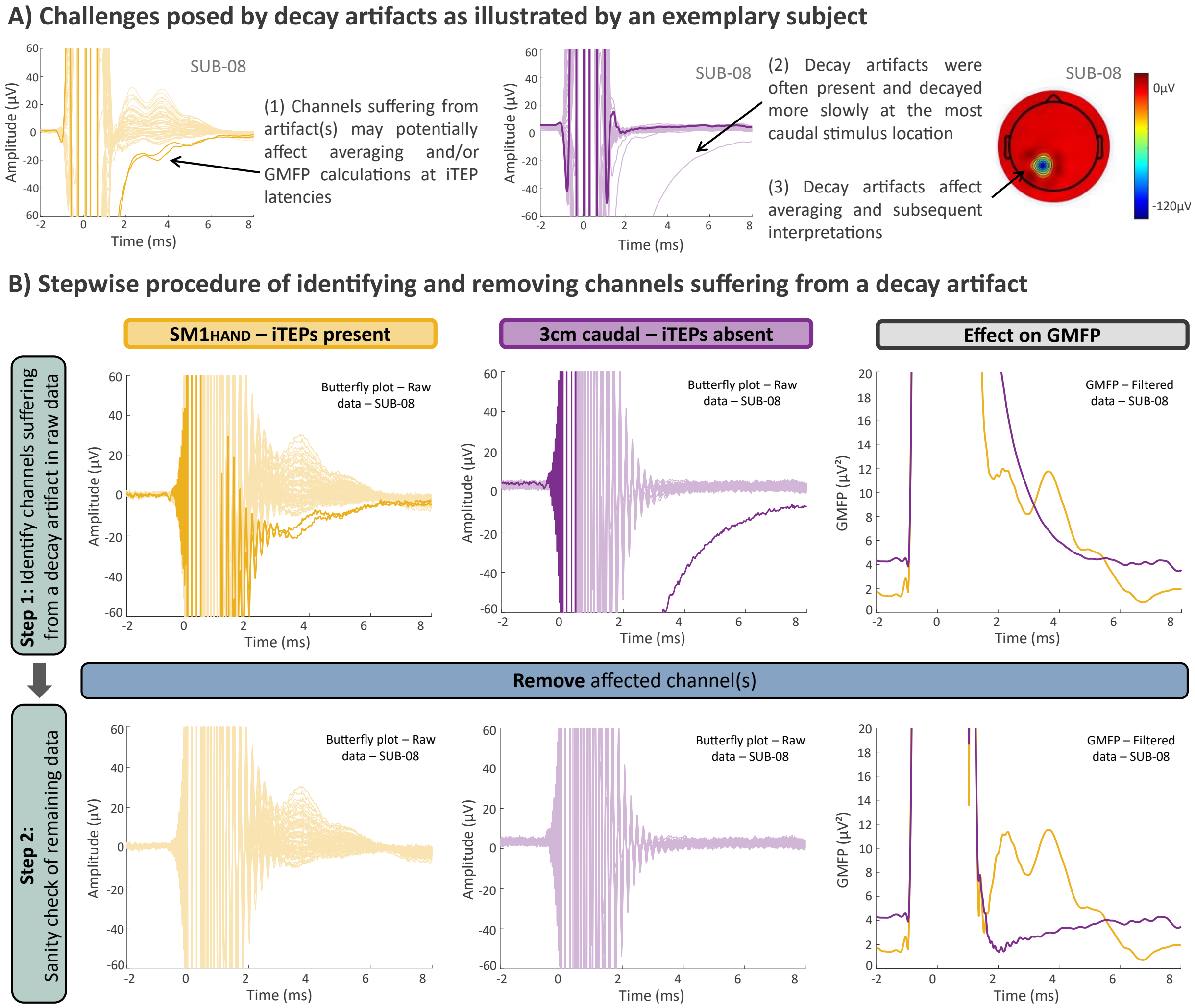


**Figure S4**. Decay artifact channels in the dataset as illustrated by an exemplary subject. **(A)** The challenges posed by decay artifacts. **(B)** Stepwise procedure for removing channels obscured by a decay artifact.

### The relationship between the onset latencies of MEPs and iTEP peaks across stimulus locations

Here, we explore the effect of stimulus location on MEP and iTEP latency. Note that rostro-caudal onset (MEPs) and peak (iTEPs) latency profiles were adjusted to the most central stimulus location. **Figure S5A** and **Table S3** indicate little change in the median adjusted onset latency of the first iTEP peak across stimulus locations. In contrast, the second and third iTEP peaks showed a systematic median increase in onset latency with increasing distance from $SM1_{HAND}$. Similarly, **Figure S5B** reveals a corresponding median increase in MEP onset latency relative to $SM1_{HAND}$. Extending this line of reasoning, **Figure S5C** explores the relationship between the differences in latency of iTEPs and MEPs over $SM1_{HAND}$. No clear relationship was observed 1 cm rostral from $SM1_{HAND}$ for the first (r = -0.19, p = 0.62) and third (r = 0.48, p = 0.52) peak, as was the case 1 cm caudal from $SM1_{HAND}$ for the first (r = 0.20, p = 0.61), second (r = 0.49, p = 0.15) and third peak. A positive but non-significant association was observed between the MEP and second iTEP peak 1 cm rostral from $SM1_{HAND}$ (r = 0.65, p = 0.058).

Overall, this exploratory analysis revealed a systematic median increase in the onset latency of later iTEP peaks, arguing in favor of their physiological nature. However, caution is warranted when interpreting these results, as the extracted onset latencies can be influenced by differences in signal-to-noise ratios or changes in the underlying slow component. In addition, although the onset of the early multi-component was around 2ms, the absolute latencies should be interpreted with caution due to the possibility of a temporal distortion caused by the filtering.

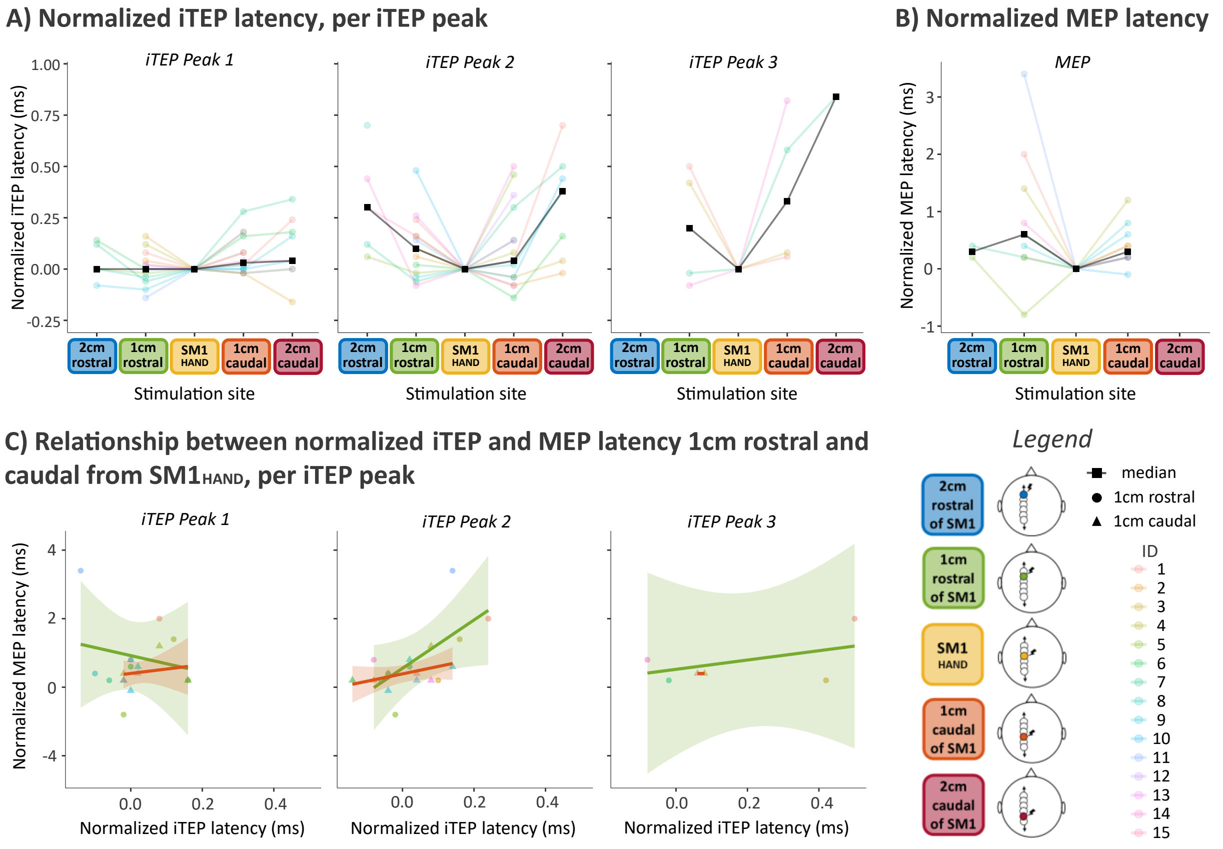


**Figure S5.** Relationship between the onset latency of immediate transcranial evoked potentials (iTEPs) and motor-evoked potentials (MEPs). **(A)** Normalized onset latency for each iTEP peak, showing a median increase in the onset latency for later iTEP peaks with increasing distance from the sensorimotor hand area ($SM1_{HAND}$). **(B)** Normalized MEP onset latency, showing a median increase in MEP onset latency moving away from $SM1_{HAND}$. **(C)** Relationship between the normalized iTEP and MEP latencies 1cm from $SM1_{HAND}$, showing that an increase in iTEP peak 2 onset latency is accompanied by a corresponding increase in MEP latency.
